# Photosynthetic CO_2_ response characteristics in canopy of *Larix principis-rupprechtii* Mayr. tree and practicability of three Models

**DOI:** 10.1101/2023.06.04.543630

**Authors:** Xuemei Ma, Chaofeng Wu, Qiang Liu, Xuanrui Huang

## Abstract

Accurately predicting the crown photosynthesis of trees is important to understand the tree growth status and carbon circle in terrestrial ecosystem. However, modeling the photosynthetic carbon dioxide (CO_2_) response curves for individual tree are still challenging due to the complex canopy structure and changeable environmental conditions. Therefore, taking 16-old year *Larix principis-rupprechtii Mayr*. as the research material, the dynamic CO_2_ response models of photosynthesis, including rectangular hyperbolic model (RHM), the non-rectangular hyperbolic model (NRHM) and the modified rectangular hyperbolic model (MRHM), were used to simulate CO_2_ response curves of the crown. The fitting accuracy of the models depend on the comparison of determinants coefficients (*R*^2^), mean square errors (*MSE*) and Akaike information criterion (*AIC*). The results showed that the mean value of *R*^2^ (*R*^*2*^=0.9939 ∼ 0.9964) of MRHM was the highest, whereas *MSE* value (*MSE*=0.2185∼0.2627) and *AIC* value (*AIC*=-13.18∼-8.03) were the lowest. The CO_2_-saturated gross photosynthetic rate (*A*_max_) and the saturation point (*C*_i_*SP*) obtained by MRHM were closest to the measured value respectively. Therefore, the MRHM fitted the CO_2_ response data well, and calculated the photosynthetic parameters directly and accurately, the fitted result showed α, *A*_max_, *C*_*i*_*SP, C*_*i*_*CP* and *R*_P_ were 0.04, 7.51 μmol·m^-2^s^-1^, 938.97 μmol·m^-2^, 67.54 μmol·m^-2^ and 0.60 μmol·m^-2^s^-1^, respectively. In addition, the difference on the photosynthetic CO_2_ response parameters values showed somewhat among different layers and orientations. In all, all data suggested that the modified rectangular hyperbolic model (MRHM) was an ideal model to fit the crown photosynthetic CO_2_ response curve of *Larix principis-rupprechtii* Mayr. These results are of great significance for parameter calibration of photosynthetic model and robust prediction of photosynthetic response in forest.

## 1 Introduction

As the largest CO_2_ fluxes in carbon, energy and other cycles in the earth system, photosynthesis can assimilate atmospheric CO_2_ and mitigate climate change, it is a critical component in the material cycle and energy flow of global terrestrial ecosystem[1-3]. CO_2_ is one of the limiting factors of photosynthesis during plant growth stage, the change of its concentrations influences the accumulation of photosynthetic products[4-7]. For the trees, the crown canopy is the most direct part for photosynthesis to response to assimilating CO_2_. Therefore, it is a major research priority to develop understanding the mechanism of leaf photosynthetic response of canopy to CO_2_ availability for improving forest productivity and ecological protection, especially for dynamic simulation of growth model and in parameterization of canopy photosynthesis.

The photosynthetic CO_2_ response curves (*A*_n_-*C*_i_ curves) elaborate the relationship between net photosynthetic rate (*A*_n_) and intercellular CO_2_ concentration (*C*_i_), the *A*_n_-*C*_i_ curve has always been one of the hotspots in plant physiology, ecology and biochemistry, it can analyze the primary productivity of vegetation and forests[8-9]. To date, in order to investigate the response of net photosynthetic rate (*A*_n_) to intercellular CO_2_ concentration (*C*_i_) of different plants, many *A*_n_-*C*_i_ curves models, including Farquhar model (1980) and its modified model [1, 10-13], the rectangular hyperbola model (RHM)[14], Michaelis-Menten model (MMM)[15], the nonrectangular hyperbola model (NRHM)[16], the exponential model (EM)[17], and the modified rectangular hyperbola model (MRHM) [18-19], the modified exponential model (MEM) [20-21], may be used to describe the the photosynthetic CO_2_ response curves (*A*_n_-*C*_i_ curves) in different plants[1], various photosynthetic parameters, such as maximum net photosynthetic rate (*A*_max_), initial carboxylation efficiency (*α*), CO_2_ saturation point (*C*_i_*SP*), CO_2_ compensation point (*C*_i_*CP*), and light respiration rate (*R*_p_), were used to assess the canopy photosynthetic capacity and other aspects of plants in different growth stage. However, RHM, MMM, NRHM, and EM were very complex, some photosynthetic parameters (such as *A*_max_ and *C*_i_*SP*) couldn’t be obtained directly from these models due to environmental changes[19, 22], and the The *C*_i_*SP* calculated by the above models was clearly underestimated, whereas *A*_max_ fitted by these models were overestimated[19]. In the contrary, due to the addition of two adjusting factors (*β* and *γ*) into the model, the MRHM can directly produce *A*_max_ and *C*_*i*_*SP*, and overcome the shortcomings of above these models, the accuracy was very higher, the results were suitable for simulating *A*_n_-*C*_i_ curves and photosynthetic parameters under different environmental conditions, it has been successfully applied to simulate CO_2_-response curves of many plants, such as *Physalis pubescens* L[23], Sa*pindus mukorossi*[9], *Alsophila spinulosa* [8], *Brassica napus* L.[24], *Nicotiana*[25].

*Larix principis-rupprechtii* Mayr (*Larch*) plays a critical role in wood production, biodiversity protection, conservation of water and soil and forest ecosystem in Northern China. Due to its complexity, little was known about the application of various dynamic canopy photosynthetic CO_2_-response models in *Larch*, and the fitting effects and differences of these models on CO_2_-response remains largely elusive. Therefore, the determinant coefficients (*R*^2^), mean square error (*MSE*), and Akaike information criterion (*AIC*) were used to evaluated the performance of three types of CO_2_-response models (such as RHM, NRHM, and MRHM) in 16-years-old *Larch*. plantation. The specific objectives of the study are to select an optimal *A*_*n*_*-C*_i_ curve model for fitting the *A*_*n*_*-C*_i_ curves of *Larch*, and to explore the relationships between the parameters of the optimal *A*_*n*_*-C*_i_ curve model and needle vertical and horizontal directions. The results will provide a theoretical guidance for further exploring the spatial heterogeneity of carbon sequestration capacity and accurately estimating photosynthetic characteristics of *Larch* needles at the crown level.

## 2 Materials and methods

### 2.1 Site description and sample tree selection

This study was conducted in the Saihanba Forest Farm of Chengde City, Hebei Province, northern China (42°02′ ∼ 42°36′N, 116°51′ ∼ 117°39′E, 1538 m a.s.l.) where contains an established plantation of *Larix principis-rupprechti* with the total area of 93,333 hm^2^. The region lies in a cold temperate semi-arid and semi humid continental monsoon climate zone in which most rainfall events occur usually between June and August, it has a mean annual precipitation of 452.6 mm, mean annual temperature of -1.5°C, evaporation of 230 mm, sunshine of 2368 hours, and mean frost-free period of 60 days, respectively. The rate of forest coverage is approximately 82.6%. Three sampling plots, each with dimensions of 20 m×30 m, were established within 16-year-old *Larix principis-rupprechtii* Mayr plantations. The average tree height and diameter at breast height of five sample trees with three plots were 12.30 m and 11.99 cm. According to previous studies[26-27], the crowns of five sample trees were divided respectively into three vertical layers (upper layer (UP), middle layer (MD), and lower layer (LW)) of equal divisions with the trisection of crown depth (the distance from the top of the tree to the bottom of its live canopy), and each layer was further divided into four orientations-east, south, west and north **(Fig. 1)[28]**.

**Fig 1.**
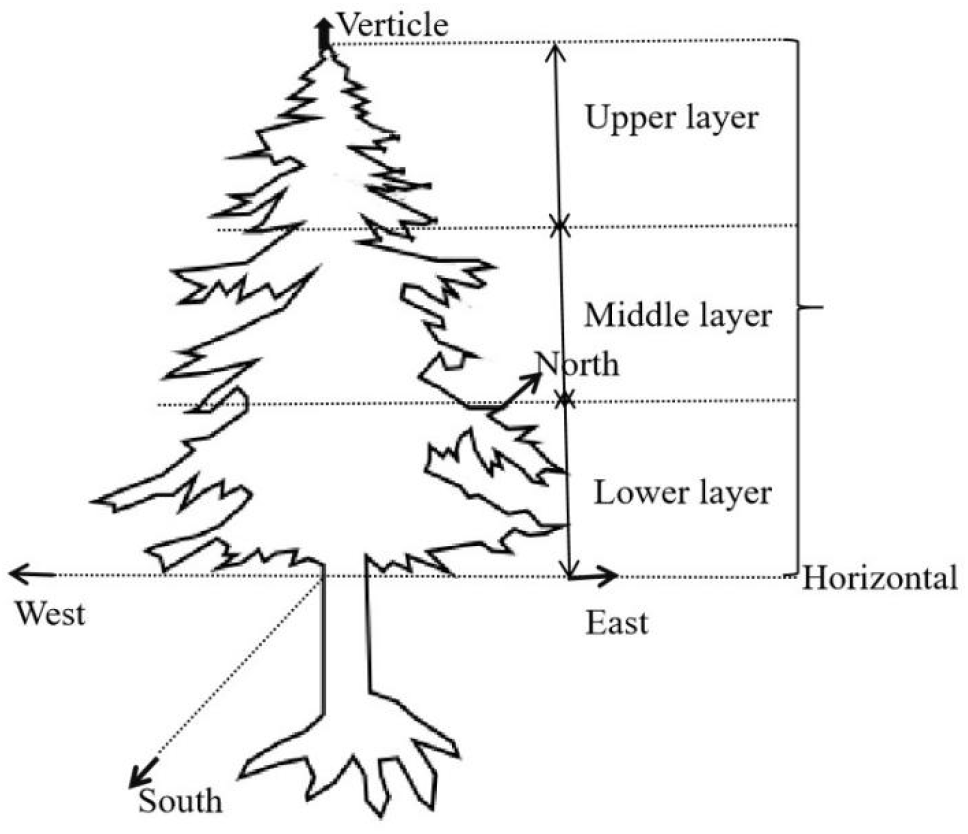
Sketch map of the canopy divisions. Upper layer, middle layer and lower layer represent three equal divisions of canopy depth in the vertical direction. East, south, west and north represent four orientations of canopy in the horizontal direction**[29]**.

### 2.2 Determination of CO_2_-response curve

The CO_2_-response curves (*A*_n_-*C*_i_ curves) for different needle positions were determined by a portable photosynthetic gas exchange system equipped with a 2 × 3 cm LED Light Source (LI-COR6400, LI-COR Inc., Lincoln, Nebraska, USA). Photosynthetic photon flux density was held at 1200 µmol m^-2^ s^-1^, temperature of leaf chamber of 24-26°C, relative humidity of 30-40% during the determination of *A*-*C*_i_ curves. Measurements of the response curves of photosynthesis to intercellular CO_2_ concentration (*A*_n_-*C*_i_) started at external CO_2_ concentration of 400 µmol mol^-1^, then decreased stepwise to 50 µmol mol^-1^ and increased stepwise to 2000 µmol mol^-1^, resulting in a sequence 400, 300, 200, 100, 50, 0, 150, 250, 400, 600,1000, 1500 and 2000 µmol mol^-1^. The experiment were conducted between 8:00 and 16:00 under cloud-free skies, the measurement data for the *A*_n_-*C*_i_ curves were obtained from June to August of 2020 and 2022. These methods were described previously[26-27].

### 2.3 Photosynthesis CO_2_-response (*A*_n_/*C*_i_) curve-fitting model

As the most frequently model for fitting *A*_n_-*C*_i_ curves, the rectangular hyperbola model (RHM), the nonrectangular hyperbola model (NRHM), and the modified rectangular hyperbola model (MRHM) were well described in the literature[19, 30-32].

**The rectangular hyperbola model (RHM)** was represented to the following form :

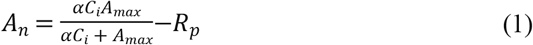

Where: *A*_n_, μmol m^-2^ s^-1^, is the photosynthetic capacity; *C*_i_, μmol·mol^-1^, is the intercellular CO_2_ concentration; *α* is initial carboxylation efficiency (dimensionless) when *C*_i_=0 μmol m^-2^; *A*_max_, μmol m^-2^s^-1^, is the maximum photosynthetic capacity; *R*_p_ is light respiration rate, it is estimated from the *A*_n_/*C*_i_ curve at *C*_i_ = 0 µmol mol^-1^.

The *A*_max_ and *C*_i_*SP* cannot be calculated directly using RHM, therefore, *A*_max_ was estimated and calculated by using the nonlinear least squares method[33], *C*_i_*SP* (saturation CO_2_ point, μmol m^-2^ s^-1^) and *C*_i_*CP* (CO_2_ compensation point, μmol m^-2^ s^-1^) were expressed on the rectangular hyperbola model in equations. 2 and 3 respectively.

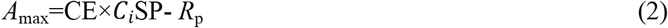

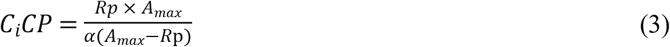

Where: CE, μmol m^-2^, is the carboxylation efficiency (CE is the slope at *C*_i_ < 200 µmol mol^-1^); *C*_i_*CP*, μmol m^-2^s^-1^, is CO_2_ compensation point; *C*_i_*SP*, μmol m^-2^s^-1^, is CO_2_ saturation point; *A*_max_ and *R*_p_ are as described above.

### Nonrectangular hyperbola model

The nonrectangular hyperbola model (NRHM) was represented to the following form:

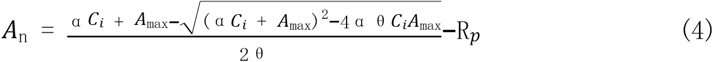

where: *θ* (0 < *θ* < 1) indicates the convexity (curvilinear angle) (dimensionless); and *A*_n_ *α, C*_*i*_, *A*_max,_ and *R*_p_ are as described above.

The *C*_*i*_*SP* and *C*_*i*_*CP* was calculated by formula 2, 5 respectively.

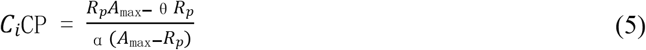

where: *C*_i_*CP, α, θ, I, A*_max,_ and *R*_p_ are as described above.

### Modified rectangular hyperbola model

The modified rectangular hyperbola model (MRHM) was represented to the following form:

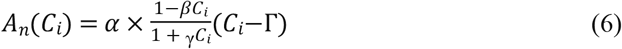

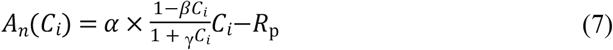

Where: Γis *C*_i_*CP, β* and *γ* are modified coefficients. *β*, μmol·m^-2^·s^-1^, represents the photoinhibition item, *γ*, μmol·m^-2^·s^-1^, represents the light saturation item, and *γ*= *α*/*P*_max_. *α, C*_i_, *R*_p_ are as described above.

The *A*_max_, *C*_i_*SP* and *C*_i_*CP* were expressed on the modified rectangular hyperbola model in equations. 8, 9, and 10 respectively:

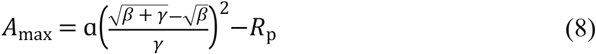

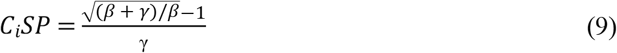

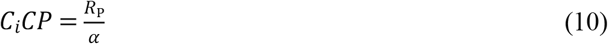

Where:*A*_max_, *α, β, γ, C*_*i*_*CP, R*_p_ are as described above.

### Model assessment and validation

The performance of the different models were assessed by determinants coefficients (*R*^2^), mean square errors (*MSE*), and Akaike information criterion (*AIC*), the best combination with the largest *R*^2^ value and smallest *MSE* and *AIC* value represented the higher fitting accuracy.

Determinants coefficients (*R*^2^) represents the fitting degree of net photosynthetic rate.

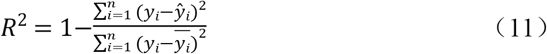

Mean square error (MSE) was the average of squared forecast errors, it is the specific value of the sum of squared errors to the number of errors.

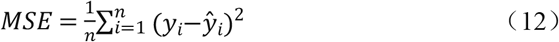

Akaike information criterion (*AIC*) is a fined technique based on in-sample fit to estimate the likelihood of a model to predict/estimate the future values.

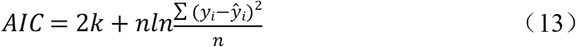

where y_i,_ ŷ_i_ and 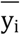 in 11, 12, and 13 equations is the measured value, the fitted value and the mean of the measured values, respectively *n* is the number of observations. *k* is the number of estimated parameters[34].

### Statistical analysis

The photosynthetic data were expressed as mean ± standard error (SE) of five independent experiments. All statistical data analyses were performed using the software GraphPad Prism 8.0 or the SPSS 18.0 (SPSS, Chicago, IL, USA). the mean values in photosynthetic parameters were significantly different at the level of 0.05 by different lowercase letters. Error bars represent the standard of the mean values of three biological replicates.

## Result

### 3.1. Spatial and temporal distribution characteristics of *A*_n_-*C*_i_

To describe the relationships between *A*_n_ and *C*_i_, photosynthetic CO_2_-response curves (*A*_n_-*C*_i_) of *Larch* was studied. The results showed the variation of *A*_n_ for needles with increasing *C*_i_. The *A*_n_-*C*_i_ curves could be divided into three phases, *A*_n_ increased rapidly as *C*_i_ was below 300 µmol mol^-1^, then increased nonlinearly up to the maximum *A*_n_ when *C*_i_ was 300-1000 µmol mol^-1^, and decreased gradually beyond 1000 µmol mol^-1^ in the third stage. The *A*_n_-*C*_i_ curves of needles under the different canopies showed similar tendency, the *A*_n_ values were very closer in the different canopy layers when the *C*_i_ was low (*C*_i_ < 300 μmol m^-2^), the gap increased with an increasing *C*_i_, and the difference in *A*_n_ was more remarkable. At the same *C*_i_, *A*_n_ under UP needles was higher than that of the MD and LW needles, respectively, and *A*_n_ values could be ranked as: upper layer > middle layer > lower layer (Fig 2, Table S1). The results indicated that needles in the upper layer have relatively higher photosynthesis, whereas those in the lower canopy layer have relatively lower photosynthesis, they are playing mutually a critical role in the canopy photosynthesis.

**Fig 2.**
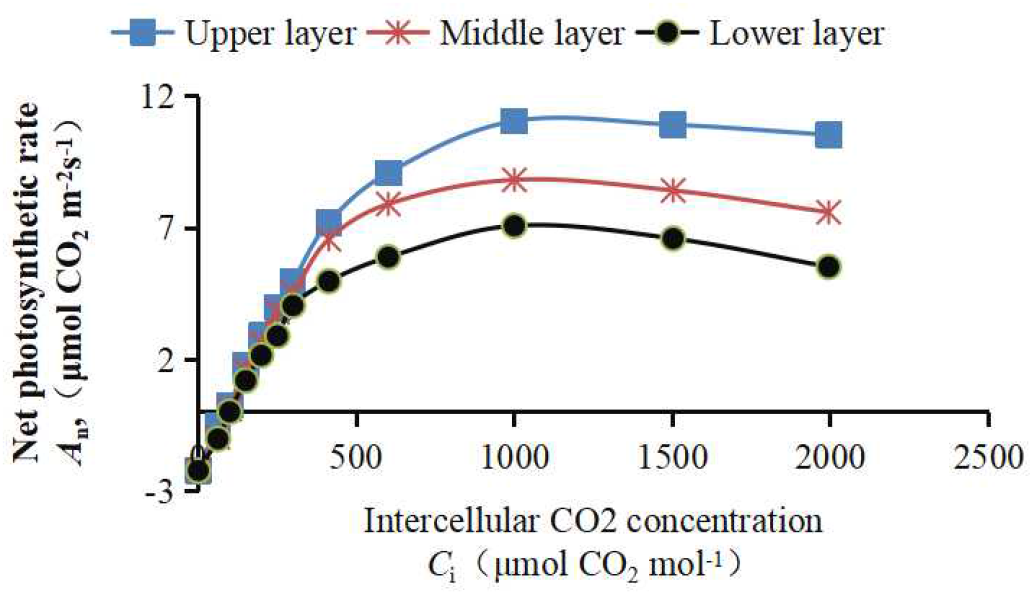
the variation of *A*_n_ for needles with increasing *C*_i_

### 3.2 Fitting and comparison of photosynthesis CO_2_ response curves in the different canopy layers and models

Fig 3 (or Table S2) showed the *A*_n_-*C*_i_ curves of needles under the different layers and different models. The *A*_n_ values fitted by the RHM, NRHM and MRHM were very close to the measured values actually when the *C*_i_ was low (*C*_i_ <300 μmol m^-2^), the *A*_n_ values simulated by three models were slightly greater than measured values when *C*_i_ values > 300 μmol m^-2^, and the pattern of *A*_n_*-C*_i_ curves simulated by three models showed similar tendency, as *C*_i_ was above 1000 μmol m^-2^, the *A*_n_ value simulated by the MRHM remained stable or decreased slowly, whereas the *A*_n_ value simulated by the RHM and NRHM increase slowly. The mean value of *R*^2^ (range from 0.9939 to 0.9964) of the MRHM model was the highest among the three models, and *MSE* value (*MSE* range from 0.2185 to 0.2627) and *AIC* value (range from-13.18 to -8.03 respective) of the MRHM were significantly smaller than those of other two models in UP, MD and LW respectively (Table. 1). Therefore, MRHM was superior to other two models in describing the photosynthetic CO_2_ response curve.

**Fig 3.**
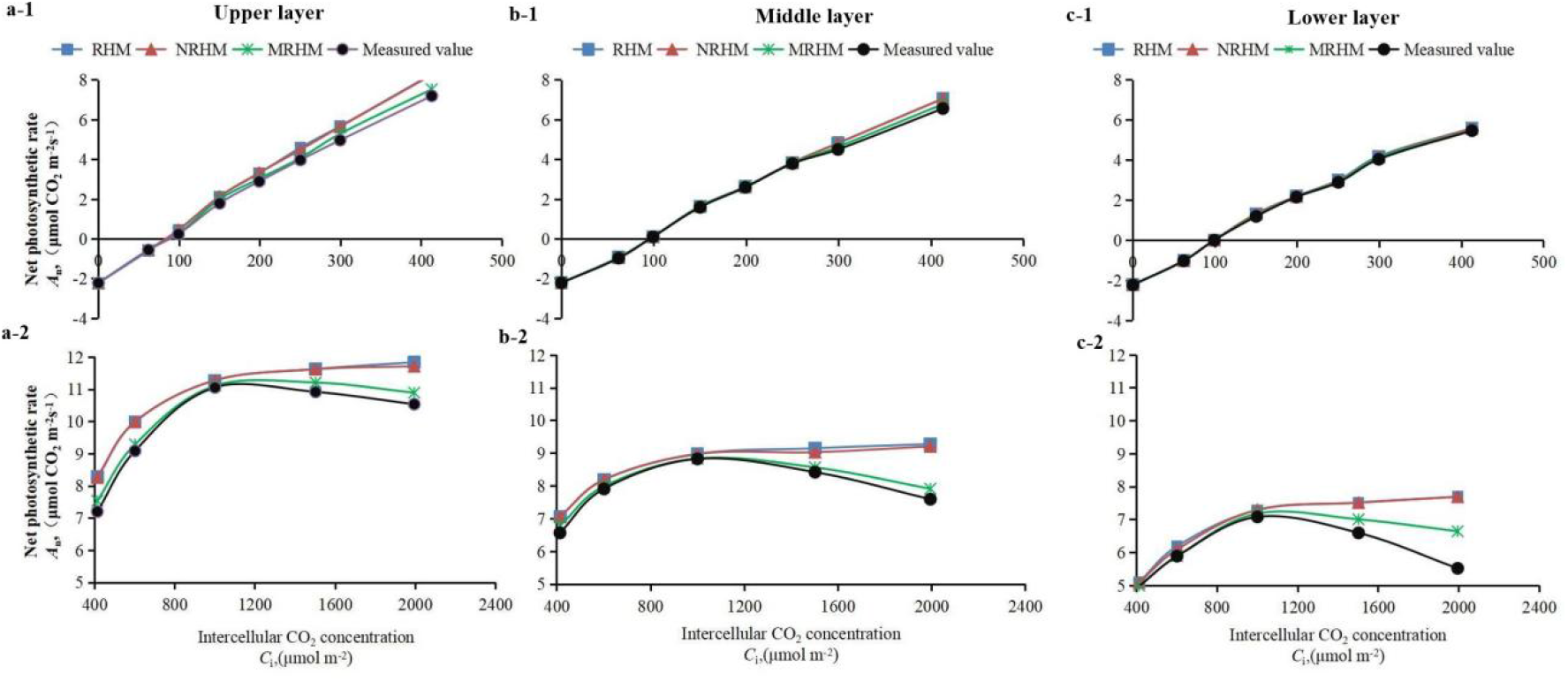
Comparison of the measured values and fitted values by three models of net photosynthetic CO_2_ response curves in different canopy layers on the RHM (a-1,a-2), NRHM (b-1,b-2), and MRHM (c-1,c-2).

**Table 1.** Fitting accuracy of different *A*_n_-*C*_i_.

### 3.3 Fitting and comparison of photosynthesis-CO_2_ response curves in the different direction

Fig 4 (or Table S3) showed the comparison of the measured and fitted values of *A*_n_ based on the MRHM in the horizontal directions. The fitted values showed little difference from the measured values actually when the *C*_i_ was low (*C*_i_ <300 μmol m^-2^), the gap increased with an increasing *C*_i_ in the different directions. *A*_n_ under FE (or ME) needles was slightly higher than that of other three directions, respectively, and *A*_n_ values were shown as: FE (or ME) > FS(or MS) > FW(or MW) > FN(or MN), the pattern of *A*_n_*-C*_i_ curves showed similar trend in the different orientations. the MRHM could fit well the variation on curves of *A*_n_ with the change of *C*_i._ In addition, MRHM was superior to other two models in each orientation (Table. 1).

**Fig 4.**
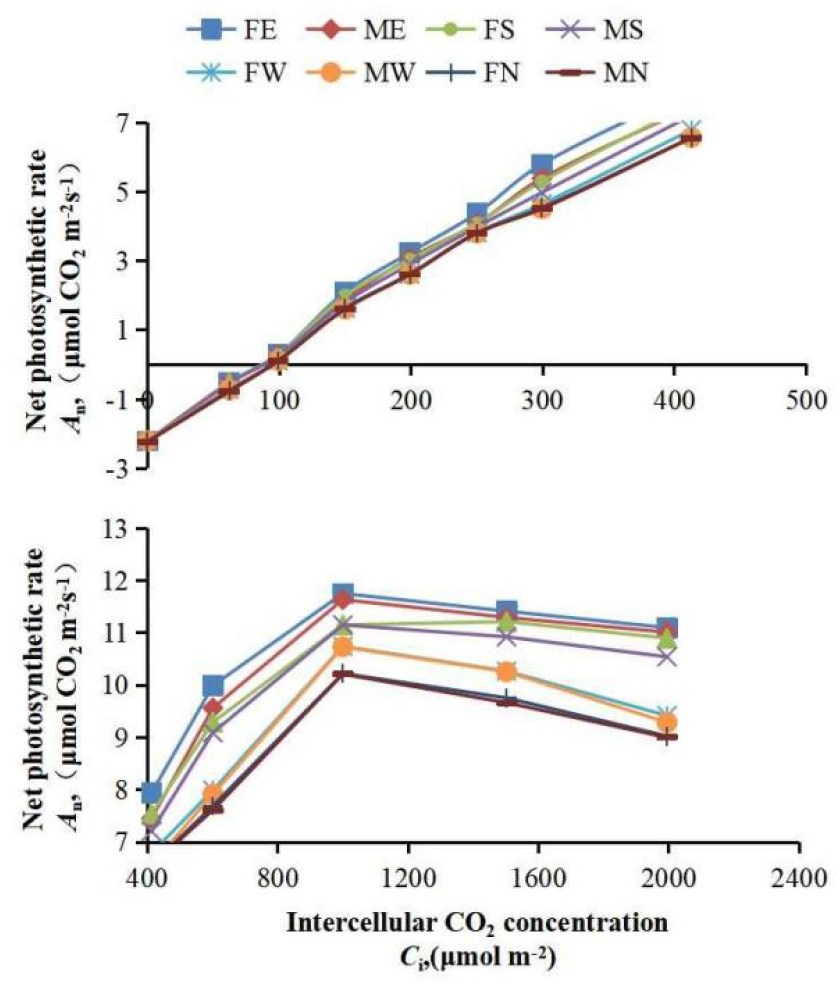
Comparison of the measured values and fitted values of net photosynthetic CO_2_ response curves in different directions on the MRHM. The letter “E”,”S”,”W”,”N” indicates the different direction of east, south, west and north, and “F”, “M” indicates the fitted value and measured value.

## 3.4 Fitting analysis of the photosynthetic parameters based on the models

It is very important to study the fitting effect of different models on photosynthetic parameters of needles. The *A*_max_ values calculated from the RHM and NRHM were significantly higher than the measured values in the upper layer, whereas the *C*_i_*SP* values calculated from the RHM and NRHM were less than the measured values in each layer. On the contrary, the fitted values of photosynthetic parameters by MRHM were very close to the measured values in different layers. In addition, the *A*_max,_ and *R*_p_ values were significant difference between MRHM and the other two models in the upper layer, while the *a*, and *C*_i_*CP* values were no significant difference among three models, the *C*_i_*SP* was significant difference between MRHM and the other two models in the different layers (Fig 5, Table S4). Therefore, MRHM could fit the photosynthetic response curve well, and better explain the saturation phenomenon of photosynthesis to CO_2_ concentrations.

**Fig 5.**
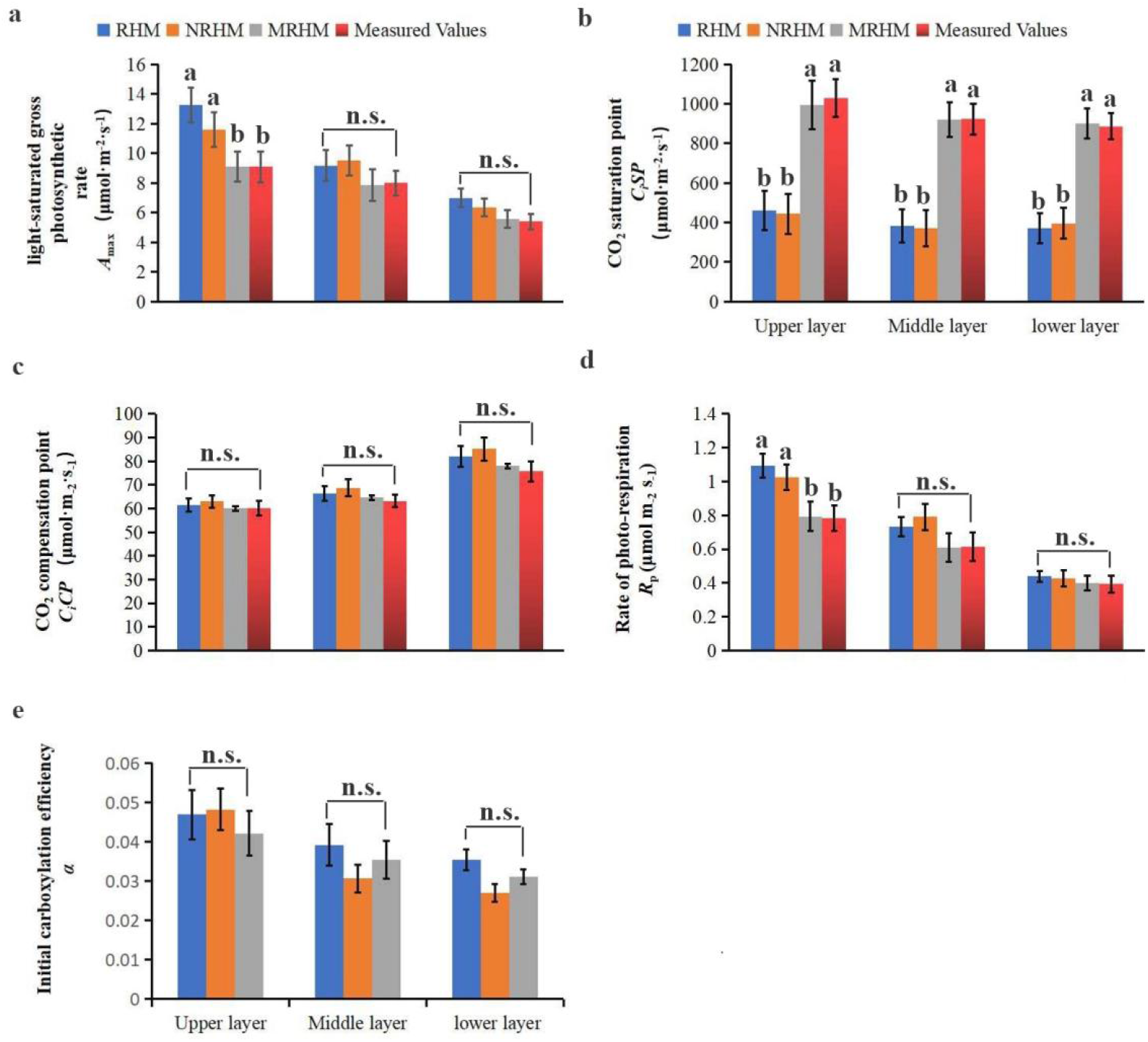
Comparison of the measured values and fitted values of photosynthetic parameters in different canopy layers on the RHM, NRHM and MRHM. Different lower case letters shows significant difference at 0.05 level, n.s. indicates no significant difference. Error bars represent the standard of the mean values of three biological replicates.

### 3.4 Comparison of the photosynthetic parameters in the different canopy layer

Fig 6 (or Table S5) showed the comparison of photosynthetic parameters of the different canopy layers. Taking the measured value and(or) the fitted value by MRHM for example, the values of *A*_max_ and *R*_p_ in the upper layer were significantly higher than that in the lower layer, and showed no difference from in the middle layer, while the value of *C*_i_*CP* in the upper was significantly lower than that in the lower layer, and showed no difference from in the middle layer. The values of *C*_i_*SP* showed no difference in different canopy layers, the difference of *a* was consistent with *A*_max_ in the vertical direction. In addition, the significant differences of the measured values of photosynthetic parameters in the different layers were consistent with the fitting values of MRHM model, and were different from those of other two models.

**Fig 6.**
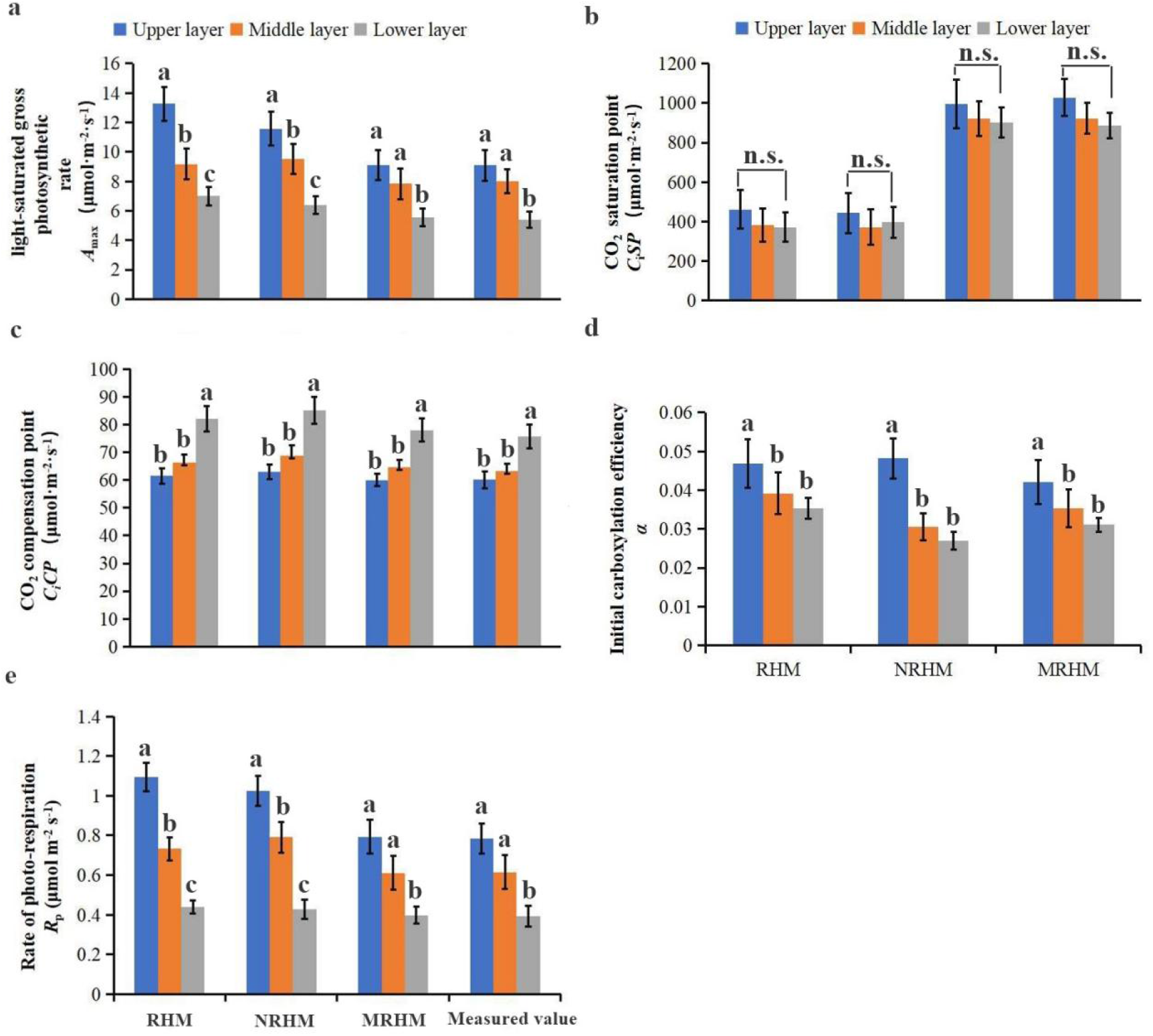
Comparison of photosynthetic parameters in different canopy layers. Different lower case letters shows significant difference at 0.05 level, n.s. indicates no significant difference. Error bars represent the standard of the mean values of three biological replicates.

## Discussion

As a critical method in elucidating the relationship between the net photosynthetic rate and CO_2_ concentrations, the *A*_n_-*C*_i_ curve can identify many photosynthetic parameters and evaluate the photosynthetic capacity of plants[35]. Therefore, choosing an appropriate CO_2_-response curve model is helpful to fit crown photosynthesis and to estimate plant photosynthetic capacity. In the study, *A*_n_ increased initially and then decreased gradually with the increase of *C*_i_ (Fig 3), which was in accordance with the study report of leaf *A*_n_-*C*_i_ curves of some plants [36]. The results implied CO_2_ absorbed by plants exceeded the needs of plants, the assimilation of the excessive CO_2_ would restricted enzymatic reaction rates in the chloroplasts and lead to delaying growth of *Larch*. In addition, the variation of *A*_n_ response to *C*_i_ maintained the state of upper layer > middle layer > lower layer at the whole growth stage, the differences of the *A*_n_-*C*_*i*_ curve among different canopy layers might be associated with leaf characteristics, higher chlorophyll a/b ratios, relative depth into crown, etc[26, 37].

The fitting of CO_2_-response curve model is a critical means to describe the response mechanism of *A*_n_ to *C*_i_ and to evaluate the photosynthetic efficiency[35]. In the study, the fitting effects of three CO_2_-response models on the *A*_n_-*C*_i_ curves of the *larch* were compared in the different layers and orientations. the fitting effects of RHM and NRHM on *A*_n_-*C*_i_ curves were better (*R*^2^>0.9) at the first stage (Fig 2), however, these two models could not fit the decline process of *A*_n_-*C*_i_ curves when *C*_i_ was above 1000 μmol m^-2^, which indicated that the application and fitting accuracy of above two models were not suitable for fitting *A*_n_-*C*_i_ of the *larch*. Compared with RHM and NRHM, the MRHM could fully fit the curve of net photosynthetic rate decreasing with the increase of CO_2_, and could reflect the actual CO_2_ saturation, it had a greater *R*^2^ and smaller *MSE* and *AIC* than other two models in simulating *A*_n_-*C*_i_ response of needles (Table. 1). In addition, some photosynthetic parameters fitted by RHM and NRHM deviated greatly from the measured values, the fitted values of photosynthetic parameters (such as α, *A*_max_, *C*_*i*_*SP, C*_*i*_*CP*) by MRHM were close to the measured values (Fig 5), which was consistent with the study of leaf *A*_n_-*C*_i_ curves of some plants at growth[5, 38-39]. Therefore, the unique structure of MRHM was more flexible in simulating different trends of *A*_n_-*C*_i_ curves [33].

The photosynthetic parameters were somewhat different in different models and canopy layers. *A*_max_, *α, C*_i_*CP* and *Rp* were significant difference in different layers (Fig 6a, 6c-6e), and was no significantly different in different models (Fig 5a, 5c-5e), while *C*_i_*SP* was no significant difference in different canopy layers (Fig 6b) and significantly different in different models (Fig 5b). Additional, the photosynthetic parameters showed no significant difference in the horizontal directions (Fig 7, Table S6). It can be seen that canopy layer was one of the main factors of affecting the photosynthetic parameters (Fig.4), the model and orientation were not an important factor that affects the photosynthetic capacity of needles. The spatial variability may be due to the comprehensive effects of genetic diversity, different producing areas and environmental factors of trees[40].

**Fig 7.**
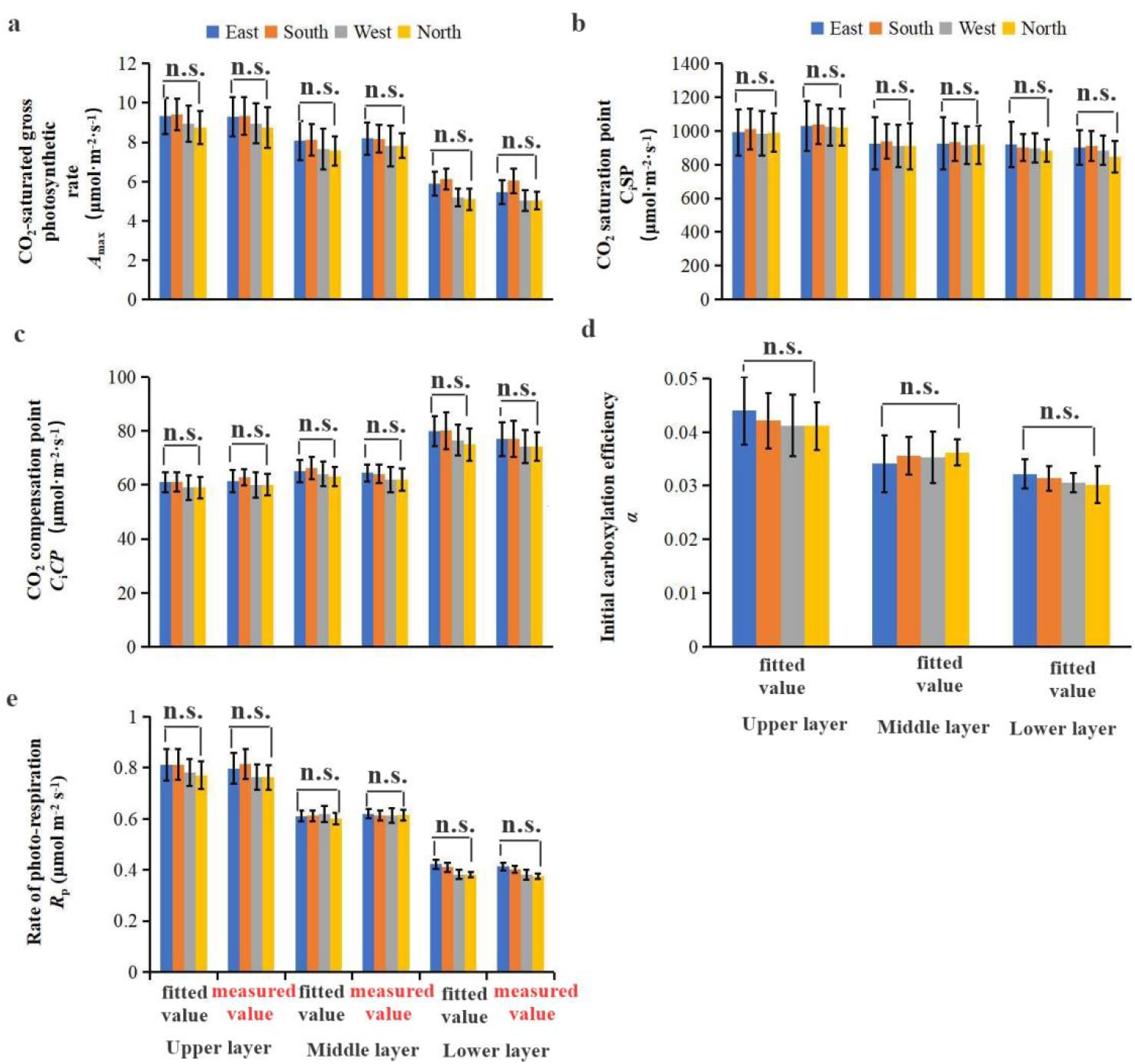
Comparison of the measured values (or fitted values) of photosynthetic parameters in different horizontal direction on the MRHM. n.s. indicates no significant difference. Error bars represent the standard of the mean values of three biological replicates.

In all, we considered the MRHM could fit well the *A*_n_-*C*_i_ curves of *larch*, and better reflect the changes of photosynthetic parameters in different canopy layers. However, the data, obtained from five sample trees of *Larix principis rupprechtii* during the growing stage in 2020 and 2021, are of some limitation for the spatial heterogeneity and complexity. Therefore, many studies are needed further to examine the relationship between *P*_n_ and *C*_i_ based on more detailed data from different sample trees of different habitat conditions, the optimal CO_2_-response curve model was selected to better understand the mechanisms of the photosynthetic physiological ecology of plants over other forest ecosystems.

## Conclusion

In the study, the *A*_n_-*C*_*i*_ curves and photosynthetic parameters were measured in different canopy layers and orientations of five planted *L. principis-rupprechtii* trees by three models at the whole growth. The results showed that the modified rectangular hyperbola model (MRHM) could simulate *A*_n_-*C*_i_ curve well and analyze the CO_2_-response data more accurately, the fitted values of photosynthetic parameters were as follows: α, *A*_max_, *C*_*i*_*SP, C*_*i*_*CP* and *R*_P_ were 0.04, 7.51 μmol·m^-2^s^-1^, 938.97 μmol·m^-2^, 67.54 μmol·m^-2^ and 0.60 μmol·m^-2^s^-1^, respectively. In addition, The photosynthesis of the *larch* has a certain spatial heterogeneity in the vertical direction. This study both helps to further explore the spatial heterogeneity of carbon sequestration capacity and provides a theoretical and practical guidance for accurately estimating the multilayered photosynthetic productivity of *Larix principis rupprechtii* plantation. We plan to extend this research to other *larch* zones and their ecosystems in northern China.

## Authorship contribution statement

Xuemei Ma: Conceptualization, Investigation, Formal analysis & Methodology. Qiang Liu: Writing -original draft, Writing - review & editing. Xuanrui Huang: Visualization, Writing - review & revision. Chaofeng Wu : Investigation, Data collection and analysis.

## Acknowledgments

The authors are grateful to Saihanba Forest Farm of Weichang County of Hebei Province for providing the material. We would like to express our appreciation to the anonymous reviewers for their constructive comments on the manuscript.

## Data Availability

All data generated and analyzed in this study are included in this published article (and its Supplementary Information Files)

## Supporting Information

**S1 Table**. the variation of *A*_n_ for needles with increasing *C*_i?_(DOC)

**S2 Table**. Comparison of the measured values and fitted values by three models of net photosynthetic CO_2_ response curves in different canopy layers on the RHM (a-1,a-2), NRHM (b-1,b-2), and MRHM (c-1,c-2). (DOC)

**S3 Table**. Comparison of the measured values and fitted values of net photosynthetic CO_2_ response curves in different directions on the MRHM. The letter “E”,”S”,”W”,”N” indicates the different direction of east, south, west and north, and “F”, “M” indicates the fitted value and measured value. (DOC)

**S4 Table**. Comparison of the measured values and fitted values of photosynthetic parameters in different canopy layers on the RHM, NRHM and MRHM. (DOC)

**S5 Table**. Comparison of photosynthetic parameters in different canopy layers. (DOC)

**S6 Table** Comparison of the measured values (or fitted values) of photosynthetic parameters in different horizontal direction on the MRHM. (DOC)

